# Nucleus accumbens and dorsal medial striatal dopamine and neural activity are essential for action sequence performance

**DOI:** 10.1101/2023.04.17.537212

**Authors:** Kurt M. Fraser, Bridget J. Chen, Patricia H. Janak

## Abstract

Separable striatal circuits have unique functions in Pavlovian and instrumental behaviors but how these roles relate to performance of sequences of actions with and without associated cues is less clear. Here we tested whether dopamine release and neural activity more generally in three striatal subdomains are necessary for performance of an action chain leading to reward delivery. Male and female Long-Evans rats were trained to press a series of three spatially-distinct levers to receive reward. We assessed the contribution of neural activity or dopamine release within each striatal subdomain when progression through the action sequence was explicitly cued and in the absence of cues. Behavior in both task variations was substantially impacted following microinfusion of the dopamine antagonist, flupenthixol, into nucleus accumbens core (NAc) or dorsomedial striatum (DMS), with impairments in sequence timing and a strong impact on motivation after NAc flupenthixol. In contrast, after pharmacological inactivation to suppress overall activity, there was minimal impact on motivation, except within the uncued task after DMS inactivation. Inactivation of both NAc and DMS impaired sequence timing and led to sequence errors in the uncued, but not cued task. There was virtually no impact of dopamine antagonism or reversible inactivation of dorsolateral striatum on either cued or uncued action sequence completion. These results highlight an essential contribution of NAc and DMS dopamine systems in motivational and performance aspects of chains of actions, whether cued or internally generated, as well as the impact of intact NAc and DMS function for correct sequence performance.

## INTRODUCTION

Reward-seeking elicited by external cues in the environment may be guided by explicit signals conveying progression towards reward or via internal models of performance. Distinct striatal subregions have been ascribed unique roles in reward-seeking with ventromedial subregions, the nucleus accumbens core (NAc) in particular, being essential for responses to Pavlovian reward-paired cues (Bornstein and Daw, 2011; Everitt and Robbins, 2013; Floresco, 2015; Collins and Saunders, 2020; de Jong et al., 2022). In contrast, more dorsal components of the striatum have been primarily ascribed the function of generating actions that lead to reward with a unique distinction between medial subregions contributing to goal-directed actions and lateral subregions contributing to habitual or automatic behaviors (Yin and Knowlton, 2006; Bornstein and Daw, 2011; Corbit and Janak, 2016; Everitt and Robbins, 2016; Lerner, 2020; Malvaez, 2020).

In the face of these functional distinctions among striatal subregions, it is of interest to consider striatal contributions to task performance that should in theory engage one or more of these region-specific processes. Here we asked whether the contributions of striatal subdivisions to reward-seeking would potentially differ when action sequences were guided by cues or executed in their absence. Because the NAc is critical for Pavlovian conditioned approach (i.e., sign-tracking) and conditioned reinforcing effects of cues, we might expect that this region contributes more strongly to cued sequence performance. In contrast, because dorsomedial striatum (DMS) is required for actions guided by knowledge of their outcome, we might expect a greater contribution of this region to uncued sequence performance. Alternatively, and perhaps more likely, normal performance in both versions of the task might require integration of the functions of these striatal subregions. For example, given evidence of its role in maintaining motivation across time delays, it is possible the NAc may contribute to sequence performance whether cued or self-paced. In addition, the DMS might be engaged to actualize action plans to achieve reward goals, whether action progression is cued or uncued. We report here that the former, not the latter, is more representative of the contributions of the striatum to the execution of action sequences, with both NAc and DMS, but not dorsolateral striatum (DLS), required for efficient performance in this task. NAc dopamine function was especially important for motivation, and the integrity of both NAc and DMS was required for sequence timing and accurate performance.

## RESULTS

### Behavioral Task and Training

We trained male and female Long-Evans rats to press a sequence of three distinct levers in order to earn a grain pellet reward (Figure 1A). We tested the effects of dopamine receptor antagonism and reversible inactivation of three striatal subregions in two variants of this three-lever sequence task. First, we assessed the impact of these manipulations when each press resulted in the cuing of the next press in the sequence to be completed by the insertion of the appropriate lever into the chamber. We then retrained rats and tested them when sequence completion was uncued as all levers remained inserted in the behavioral chamber throughout the session with only a brief cue associated with reward delivery indicating sequence completion. Rats were implanted with cannula prior to training.

**Figure 1.**
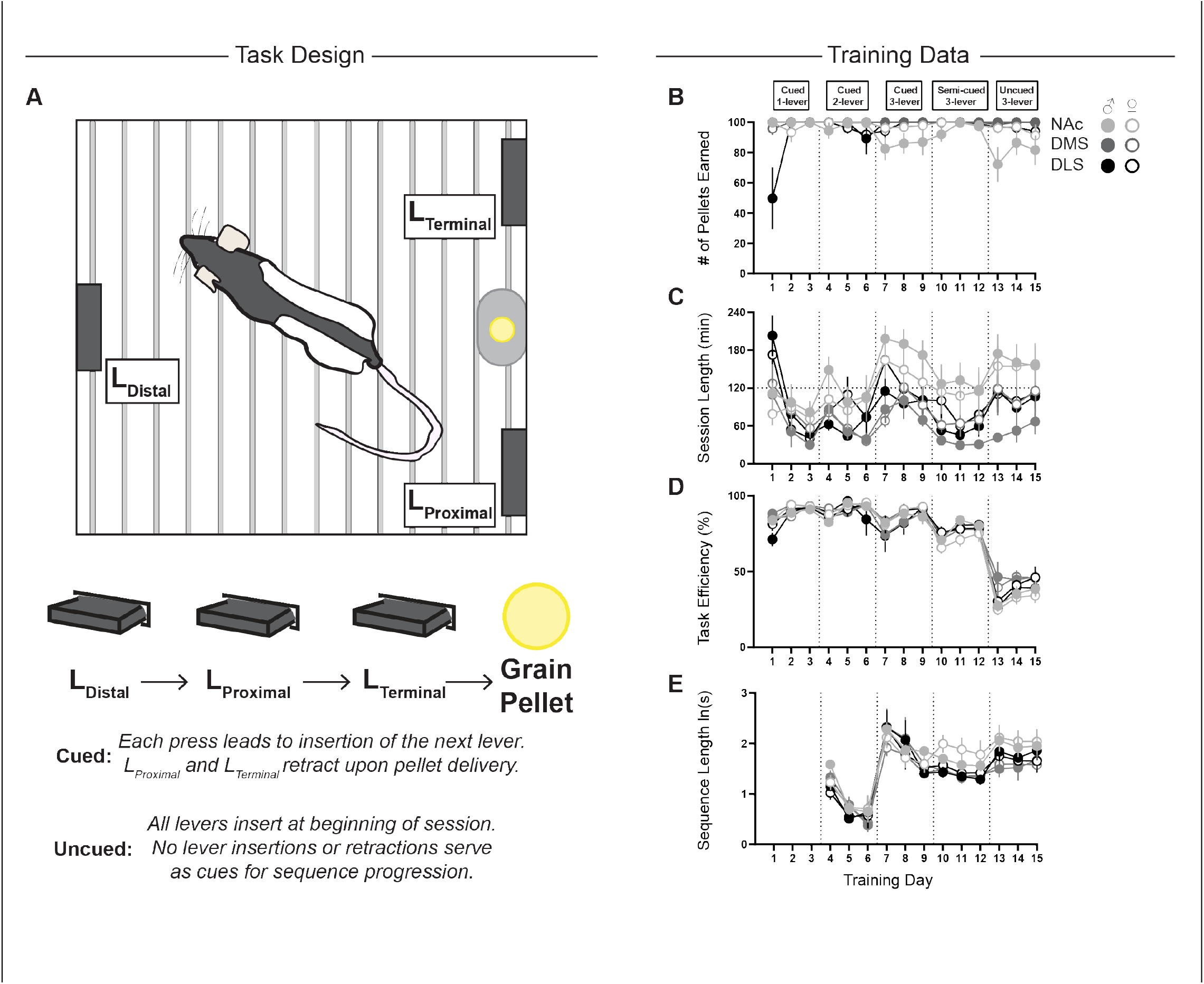
Task design and training data. **A)** Overview of general task design where rats were required to press three distinct levers in sequence to earn a 45 mg grain pellet reward. We tested rats when sequence progression was cued by the insertion of the next lever in sequence following each press and also when each lever was constantly available during a session. **B)** Rewards earned in each training session. **C)** Session duration. **D)** Efficiency of performance in the task expressed as percentage of correct to total lever presses. **E)** Average length of each sequence. For all graphs, data are mean ± SEM. Closed symbols represent male rats and open circles represent female rats. NAc, nucleus accumbens; DMS, dorsal medial striatum; DLS, dorsal lateral striatum.

There were no significant sex differences detected during training for rewards earned (Figure 1B; group by sex by day F(28,64.949)=1.463, p=0.105 ; group by day F(28,64.949)=1.728, p=0.036; sex by day F(14,64.949)=1.189, p=0.305), session completion time (Figure 1C; group by sex by day F(28,66.519)=0.678, p=0.872; group by day F(28,66.519)=2.347, p=0.002; sex by day F(14,66.519)=0.429, p=0.960), or average sequence length (Figure 1E; group by sex by day F(22,65.089)=1.049, p=0.423 ; group by day F(22,65.089)=1.797, p=0.036; sex by day F(11,65.006)=1.140, p=0.346), with the exception of efficiency of sequence performance (Figure 1D; group by sex by day F(28,50.464)=1.785, p=0.036; group by day F(28,50.464)=1.485, p=0.110; sex by day F(14,50.466)=0.429, p=0.086). In the case of performance efficiency, it is important to note that we did not observe significant effects of sex within each group on the three training days (7-9 or 13-15, respectively) that occurred prior to each series of tests (effects of sex within each group p>0.05).

When we examined significant group by day interactions in the training data these primarily detected significant differences between rats with NAc cannulae and either DMS or DLS cannulated rats on select days across measures, although the general rate of learning and responding was consistent within each of these groups (significant effects of day within each group p<0.05). These differences across groups may have arisen from basal differences in each group prior to implantation of cannula or from the cannulation surgery/presence of cannula. Nonetheless, it is apparent all groups acquired the task and performance was equivalent between male and female rats. We next tested the effects of dopamine antagonism and reversible inactivation in these three distinct striatal subregions (Figure 2).

**Figure 2.**
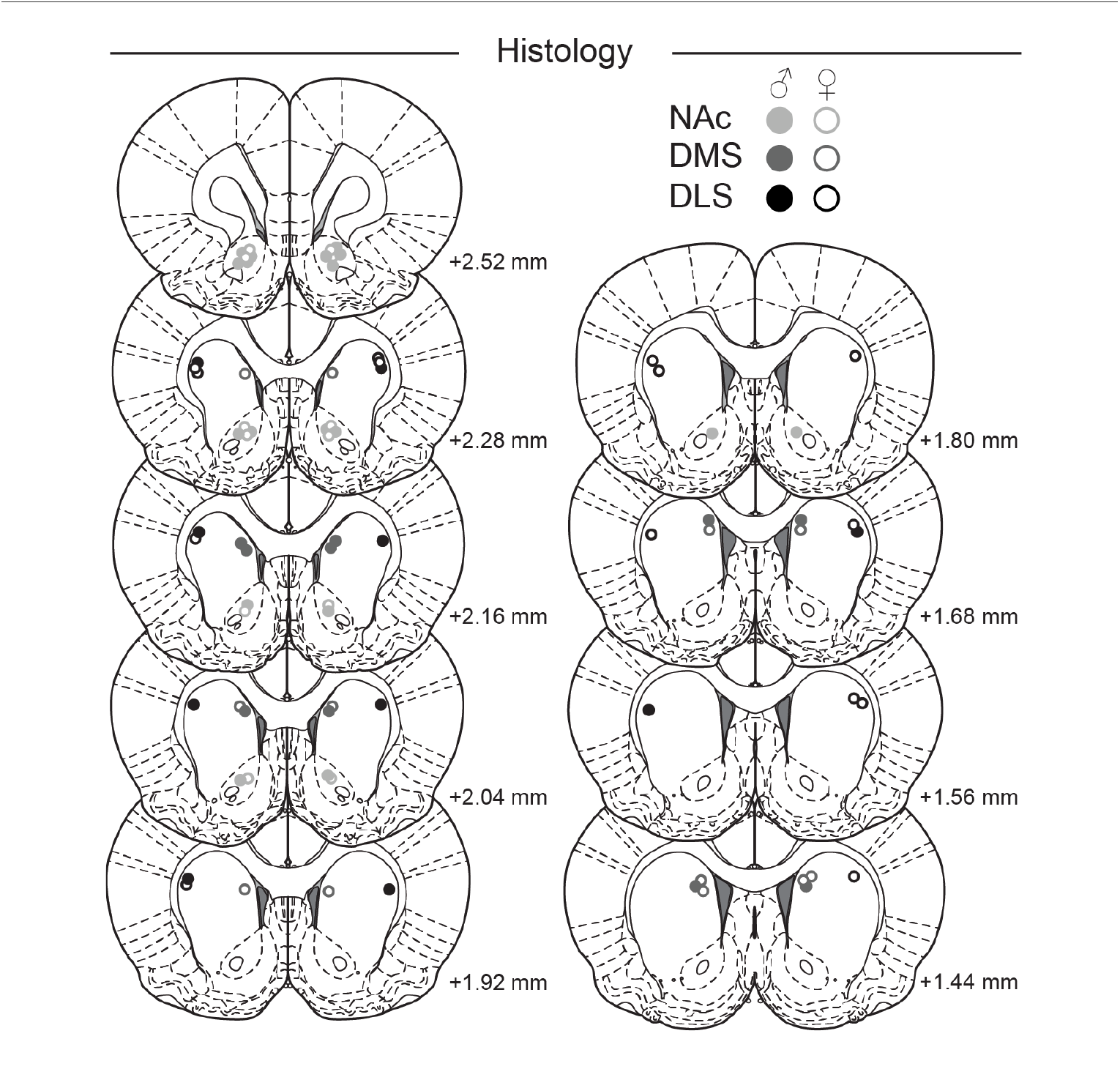
Histological verification of microinfuser tip location. Reconstruction of the final placement of microinfuser tips with location relative to bregma indicated. Closed symbols represent male rats and open circles represent female rats. Atlas sections adapted from (Paxinos and Watson, 2007).

### Nucleus Accumbens Core Dopamine and Neural Activity Differentially Control Cued and Uncued Sequence Completion

When sequence progress was cued by the insertion of each lever, blockade of dopamine receptors in the NAc via microinfusion of flupenthixol decreased the number of rewards earned out of a maximum of 100 (Figure 3A; F(1.631,22.83)=22.06, p<0.0001; posthoc p<0.0001) and increased session duration (Figure 3B; F(1.989,27.85)=7.873, p=0.0020; posthoc p=0.0089), while there was no comparable impact of reversible inactivation (post-hocs p>0.05). Despite the lack of significant impact on motivation to earn rewards, reversible inactivation did reduce efficiency (Figure 3C; F(1.448,19.55)=5.402, p=0.0209; posthoc p=0.0176), an effect that also held true for dopamine receptor antagonism (p=0.0064). There was no effect of either microinfusion treatment on the interval between sequences (Figure 3D; F(1.185,15.41)=3.623, p=0.0703). Although the mean overall sequence length was not statistically significantly altered (Figure 3E; F(1.096,14.79)=4.276, p=0.0537),. both reversible inactivation and dopamine antagonism in the NAc selectively lengthened the first inter-reponse interval, between LD and LP (Figure 3F; interaction of treatment and sequence segment F(2,26)=4.815, p=0.0166; treatment F(2,28)=3.922, p=0.0315; posthocs p<0.05). Neither treatment significantly affected the number of errors when the sequence was cued (Figure 3G; interaction of treatment and error type F(2,28)=1.085, p=0.3517; treatment F(2,28)=1.221, p=0.3100). Thus, NAc dopamine is essential for continued performance of a cued sequence of lever presses, as seen by deficits in completing all trials, while suppression of NAc activity more broadly does not impact motivation to complete the session. Both treatments proportionally increase extra lever press responding and selected inter-response intervals.

**Figure 3.**
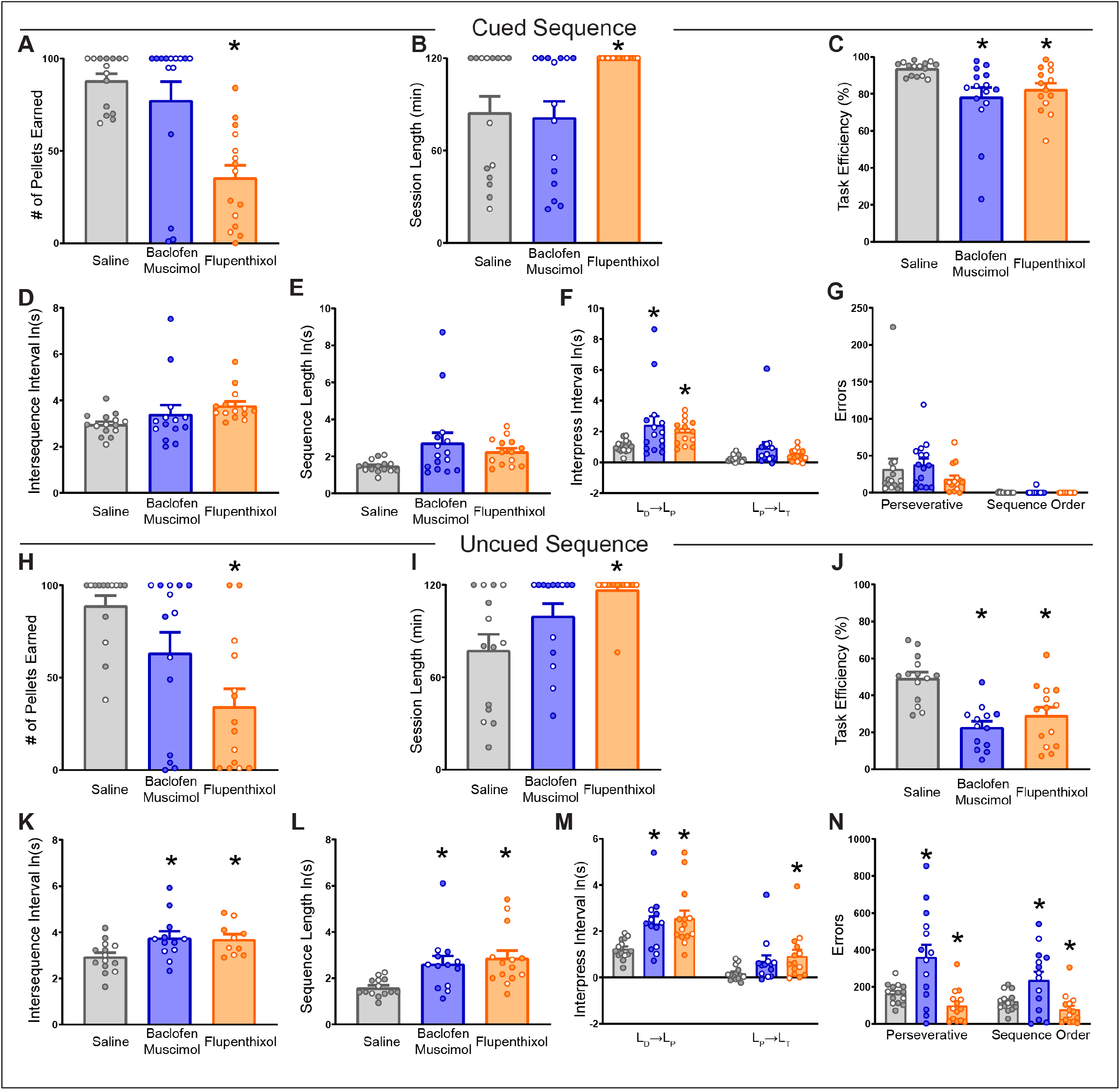
Nucleus accumbens core dopamine receptor blockade and suppression of neural activity affect multiple aspects of sequence performance. **A)** Rewards earned in each test session. **B)** Session duration. **C)** Efficiency of performance in the task expressed as percentage of correct to total lever presses. **D)** Average time to initiate a new sequence following the last reward delivery. **E)** Average length of each sequence. **F)** Average length of the time between each sequence segment from either LD to LP or LP to LT. **G)** Total number of perseverative errors and sequence order errors at test. **H-N)** same as **A-F** but data represent test sessions when all levers were continuously available and no cues served to guide sequence progress. Data are mean ± SEM. Closed symbols represent male rats and open circles represent female rats. * p<0.05.

We then retrained and tested these rats when performance and progression was uncued with all levers continuously available throughout a session. Similar to when progress was cued, dopamine receptor antagonism in the NAc reduced rewards earned (Figure 3H; F(1.713,22.27)=12.87, p=0.0003;posthocp<0.0001)andincreasedsession completion time (Figure 3I; F(1.631,21.20)=6.598, p=0.0086; posthoc p=0.0027). There was no significant impact of inactivation on either of these measures suggesting a more modest effect of NAc inactivation on motivation to earn rewards (posthoc p’s>0.05). Both treatments also reduced task efficiency (Figure 3J; F(1.915,22.68)=17.64, p<0.0001; posthocs p<0.001). Both reversible inactivation and dopamine blockade increased the time it took for rats to initiate a new sequence (Figure 3K; F(1.501,15.01)=6.656, p=0.0128; posthocs p<0.05) and the mean sequence length (Figure 3L; F(1.961,24.51)=9.358, p=0.001; posthocs p<0.05) and this primarily resulted from a lengthening of the first segment of the sequence (Figure 3M; treatment F(2,26)=8.226, p=0.0017; posthocs p<0.01). Despite generally similar treatment effects on performance, inactivation of the NAc additionally increased perseverative errors (Figure 3N; treatment F(2,26)=121.10, p=0.0003; interaction of treatment and error type F(2,26)=10.03, p=0.0006; posthoc p<0.0001) and sequence order errors (p<0.0001), while NAc flupenthixol reduced perseverative (p=0.0012) and sequence order errors (p=0.0282).

Taken together these results suggest a critical role for NAc dopamine in both the motivation to perform and the execution of action sequences irrespective of the presence of cues indicating progress towards reward. Of note, flupenthixol did not increase sequence errors. Interestingly, while pharmacological inactivation of the core also impacted execution, this temporary reduction in neural activity also produced errors in responding, while leaving overall motivation to complete the session less affected. Together these findings indicate roles for NAc in maintaining motivation to pursue reward, as well as in the appropriate selection of the specific motor actions necessary to maintain fluid progress throughout a sequence, limit errors, and ultimately refine behavioral output.

### Manipulation of Dorsomedial Striatal Dopamine and Neural Activity Also Differentially Alters Cued and Uncued Sequence Completion

We next tested, in separate rats, whether dopamine release or neural activity in DMS was required for sequence performance. When sequence progression was cued by the insertion of the appropriate lever to press, microinfusion of flupenthixol into the DMS reduced the number of rewards earned (Figure 4A; F(1.691,16.91)=5.821, p=0.0150; posthoc p=0.0069) and in turn increased session completion time (Figure 4B; F(1.528,15.28)=13.73, p=0.0008; posthoc p=0.0001). There was no impact of reversible inactivation on either rewards earned (posthoc p=0.2848) or session completion time (posthoc p=0.9759), suggesting that DMS neural activity is not necessary for motivation to earn rewards as indicated by completion of all trials when sequence progression is cued. There was an overall impact of both drug treatments within the DMS on task efficiency (Figure 4C; main effect of treatment F(1.694,16.94)=4.386, p=0.0343) and the time to initiate a new sequence following reward (Figure 4D; F(1.694,16.94)=3.874, p=0.0471), yet no posthoc comparisons were statistically significant compared to saline (p’s>0.05). In agreement with flupenthixol reducing rewards earned, the average sequence length was significantly extended following this treatment (Figure 4E; F(1.817,18.17)=3.820, p=0.0447) but not by reversible inactivation (p=0.0703). Interestingly, when we analyzed each sequence segment separately we observed a significant effect of both drug treatments on lengthening the time to complete the first sequence segment from LD to LP (Figure 4F; interaction of treatment and sequence segment F(2,20)=5.942, p=0.0094; treatment F(2,20)=3.684, p=0.0435; posthocs p<0.001). There was no effect of either drug in the DMS on errors when sequence progression was explicitly cued (Figure 4G; interaction of treatment and error type F(2,20)=3.285, p=0.0584; treatment F(2,20)=3.245, p=0.0602).

**Figure 4.**
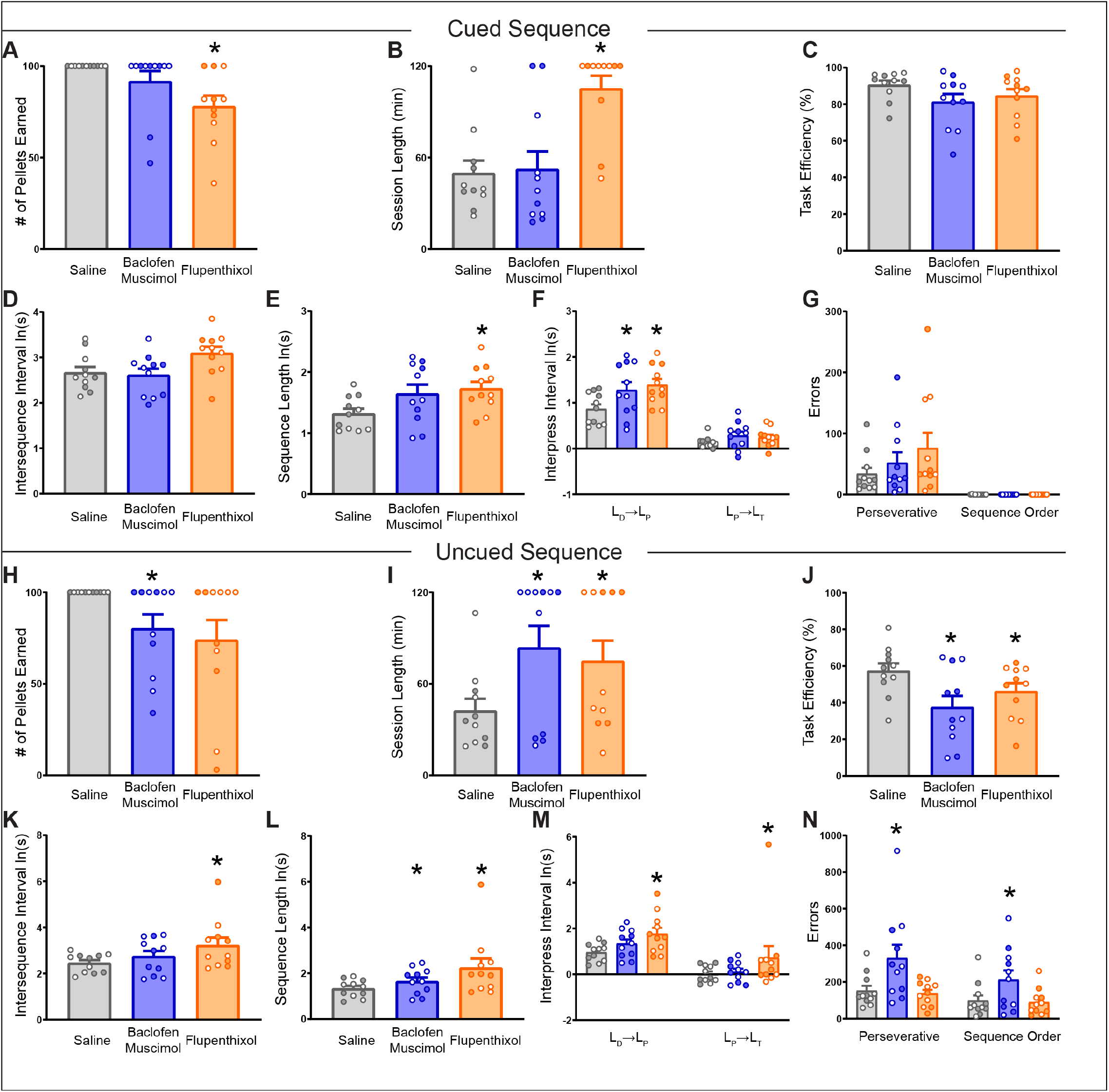
Dorsal medial striatal dopamine receptor blockade and suppression of neural activity affect multiple aspects of sequence performance. **A)** Rewards earned in each test session. **B)** Session duration. **C)** Efficiency of performance in the task expressed as percentage of correct to total lever presses. **D)** Average time to initiate a new sequence following the last reward delivery. **E)** Average length of each sequence. **F)** Average length of the time between each sequence segment from either LD to LP or LP to LT. **G)** Total number of perseverative errors and sequence order errors at test. **H-N)** same as **A-F** but data represent test sessions when all levers were continuously available and no cues served to guide sequence progress. Data are mean ± SEM. Closed symbols represent male rats and open circles represent female rats. * p<0.05.

We retested these rats when each lever remained available throughout each session, removing the ability of lever insertion to serve as cues for sequence progress. Reversible inactivation significantly reduced the number of rewards earned (Figure 4H; F(1.375,13.75)=5.135, p=0.0312; posthoc p=0.0488) and while dopamine antagonism reduced rewards earned as well, this was nonsignificant (p=0.0677). Despite this discrepancy, both dopamine receptor antagonism and reversible inactivation significantly increased session duration (Figure 4I; F(1.511,15.11)=5.605, p=0.0211; posthocs p<0.05), reduced task efficiency (Figure 4J; F(1.885,18.85)=11.30, p=0.0007; posthocs p<0.05), and increased the time it took to complete the 3-lever sequence (Figure 4L; F(1.084,10.84)=4.884, p=0.0473; posthocs p<0.05). Interestingly, flupenthixol in the DMS significantly lengthened the time it took for rats to initiate a sequence following reward (Figure 4K; F(1.489,14.89)=6.175, p=0.0164; posthoc p=0.0274), but there was no significant effect of inactivation (p=0.1259). When we analyzed the length of each sequence segment individually there was a significant effect of treatment (Figure 4M; F(2,20)=4.539, p=0.0237) reflecting an increase in the length of each segment following dopamine antagonism (p=0.0159). Finally, we considered whether each treatment altered errors during the behavioral session. Interestingly, we found that reversible inactivation of the DMS resulted in a significant increase of both perseverative and sequence order errors (Figure 4N; treatment F(2,20)=8.157, p=0.0026; posthoc p=0.0063) with no impact of dopamine antagonism on error production (p=0.9535). Collectively, DMS dopamine signaling regulates both the motivation to seek reward and the speed with which sequences are completed regardless of cuing of actions. While DMS inactivation also impacts motivation to complete the session, this appears to only hold for the uncued version of the task, with similar impacts on sequence performance speed for both cued and uncued sequences. In addition, DMS neural activity is essential for limiting the number of erroneous lever presses when sequence completion is self-paced and without inter-sequence feedback.

### Neither Dorsolateral Striatal Dopamine Signaling nor Neural Activity are Essential for Sequence Performance

In contrast to the effects observed with identical manipulations in the NAc or DMS, dopamine receptor antagonism or reversible inactivation of the DLS had minimal effects on sequence performance. We found no effect of either treatment on rewards earned (Figure 5A; F(1.120,12.32)=1.964, p=0.1865), session length (Figure 5B; F(1.799,19.78)=1.248, p=0.3051), the intersequence interval (Figure 5D; F(1.822,20.70)=0.8253, p=0.4454; posthocs p<0.05), sequence length (Figure 5E; F(1.915,21.06)=3.063, p=0.0699), or number of errors (Figure 5G; treatment F(2,22)=2.806, p=0.0821; interaction F(2,22)=2.806, p=0.0821) madewhensequenceperformancewascued.There were significant effects of flupenthixol in reducing task efficiency (Figure 5C; F(1.921,21.13)=4.565, p=0.0236; posthoc p=0.0305) and lengthening the time it took for rats to progress from the first LD press to the subsequent LP press (Figure 5F; interaction of treatment and sequence segment F(2,22)=9.403, p=0.0011; posthoc p=0.0004).

**Figure 5.**
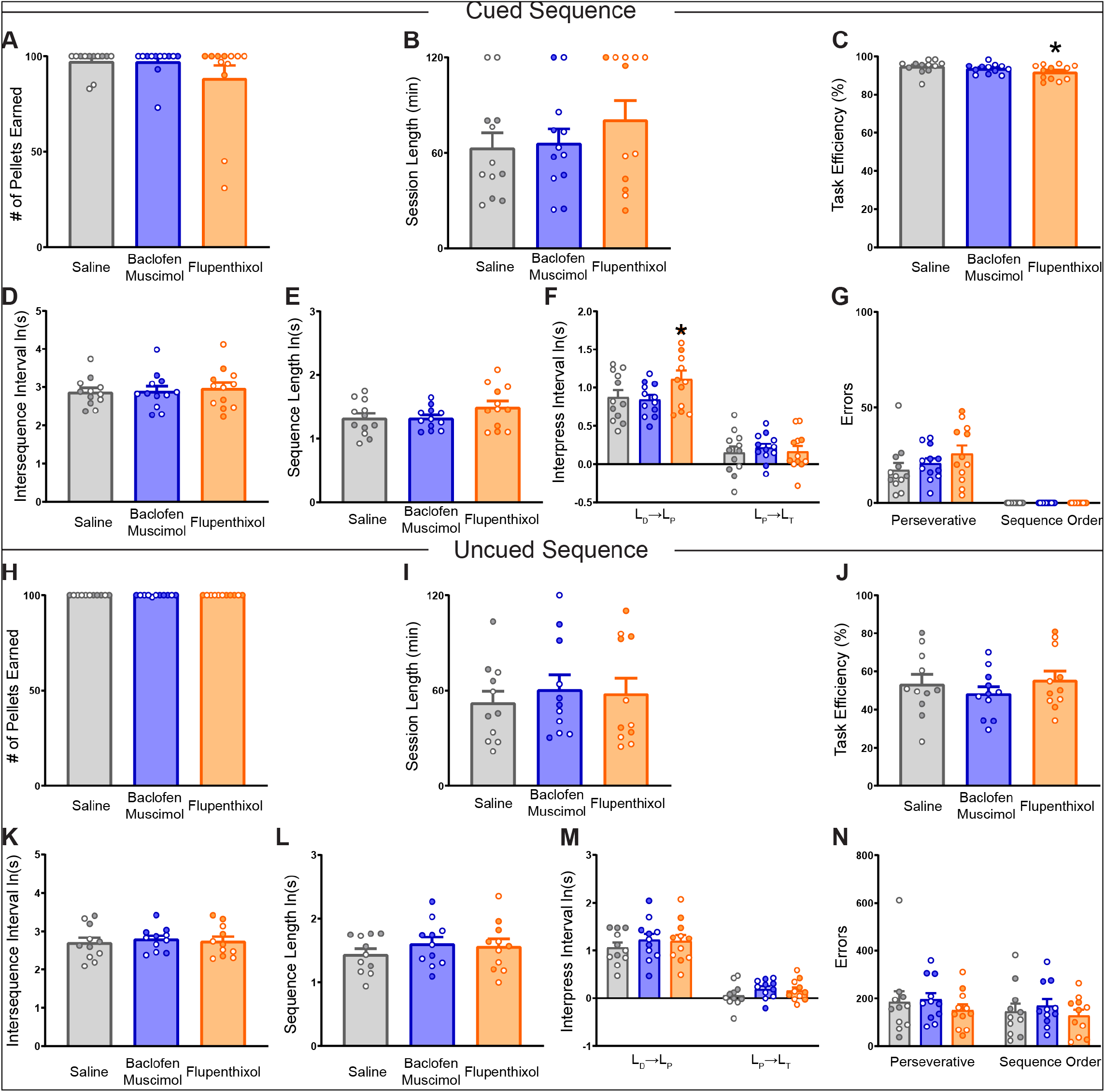
Minimal contribution of dorsal lateral striatal dopamine receptor blockade or neural activity suppression to sequence performance. **A)** Rewards earned in each test session. **B)** Session duration. **C)** Efficiency of performance in the task expressed as percentage of correct to total lever presses. **D)** Average time to initiate a new sequence following the last reward delivery. **E)** Average length of each sequence. **F)** Average length of the time between each sequence segment from either LD to LP or LP to LT. **G)** Total number of perseverative errors and sequence order errors at test. **H-N)** same as **A-F** but data represent test sessions when all levers were continuously available and no cues served to guide sequence progress. Data are mean ± SEM. Closed symbols represent male rats and open circles represent female rats. * p<0.05.

Despite a restricted impact of our treatments on sequence execution when progress was cued, we assessed if DLS was recruited to maintain sequence performance if cues were removed. Neither dopamine antagonism nor reversible inactivation within the DLS impacted sequence performance. There was no observed effect of treatment on rewards earned (Figure 5H; F(1,10)=1.000, p=0.3409), session completion time (Figure 5I; F(1.718, 17.18)=0.6112, p=0.5303), efficiency (Figure 5J; F(1.660,16.60)=1.708, p=0.2130), time between sequences (Figure 5K; F(1.764,17.64)=0.6259, p=0.5269), and sequence length (Figure 5L; F(1.568,15.68)=2.527, p=0.1206) nor an interaction or main effect of treatment on the length of each sequence segment (Figure 5M; interaction of treatment and sequence segment F(2,20)=0.0502, p=0.9512; treatment F(2,20)=2.604, p=0.0988) and errors made (Figure 5N; interaction of treatment and error type F(2,20)=0.4570, p=0.6403; treatment F(2,20)=0.9039, p=0.4209). These results collectively indicate a lack of requirement for dopamine signaling and neural activity in the DLS to sequence performance in this task irrespective of action cuing.

## DISCUSSION

Motivated pursuit of reward typically involves chains of behaviors that close the gap between an agent and reward. In some cases, cues may explicitly signal the proper completion of individual actions to help shape reward-seeking and in other instances pursuit of reward occurs in the absence of immediate feedback. We assessed here the contribution of three striatal subdivisions in variations of these forms of reward-seeking actions in rats performing a sequence of lever presses at three distinct locations to earn reward. We find dopamine release in the nucleus accumbens core (NAc) and the dorsomedial striatum (DMS) is essential for maintaining motivation across the session to work for reward whether performance was cued or not, with a greater magnitude of effect after flupenthixol microinfusion in the accumbens. In addition, dopamine receptor blockade in both the NAc and DMS impacted measures of sequence performance and these effects were more noticeable in the uncued version of the task. Thus, we conclude dopamine signaling in both these striatal subregions is normally required for appropriate performance of a chain of instrumental responses, whether those responses are cued or self-initiated. In contrast, there was little impact of reversible silencing of neural activity in the NAC and DMS on measures of motivation, but there were consistent effects on sequence execution as reflected by increases both in time to complete a sequence once initiated and in extra, non-essential lever presses. The effects observed after both NAc and DMS inactivation were particularly profound during self-paced sequence performance in the absence of cues to provide feedback. In addition, in the uncued version of the task, NAc and DMS inactivation increased the number of erroneous lever presses, both perseverative and those reflecting errors in sequence memory. Surprisingly, we found that there were negligible effects of dopamine antagonism or reversible inactivation of the dorsolateral striatum (DLS) in sequence performance irrespective of the presence or absence of cues. These results suggest dissociations in the contributions these distinct striatal subdivisions to motivation, correct sequence generation, and action execution.

In both the cued and uncued versions of the task, dopamine antagonism within the core region of the accumbens strongly reduced task participation, suggesting an effect on motivation to respond in agreement with prior findings within instrumental tasks (e.g., (Cousins et al., 1994; Fischbach-Weiss et al., 2017). Interestingly, DMS flupenthixol infusion had impacts on motivation as well, as indicated by increases in session length, although we note these appear smaller in effect than those observed with nucleus accumbens core manipulations. While dopamine receptor blockade in both the NAc and DMS generally decreased the number of rewards acquired, there was no increase in performance errors, in contrast to overall suppression of activity in these areas. Indeed, after flupenthixol infusion in the NAc, rats showed fewer errors, both perseverative and sequence. We also found that NAc and DMS dopamine blockade lengthened the time it took for rats to initiate a new sequence, in agreement with reports of dopamine neuron activity in the ventral tegmental area and substantia nigra facilitating action sequence initiation (Fischbach-Weiss et al., 2017; da Silva et al., 2018). In addition, the dopamine antagonist increased time to execute the sequence, especially when it was uncued. These performance effects are in agreement with prior findings of ventral tegmental area dopamine neuron optoinhibition on initiation and execution of an instrumental sequence in mice (Fischbach-Weiss et al., 2017).

The striatal circuits studied here are embedded in a hierarchical spiraling network with midbrain dopamine neurons and a training-related progression of dopamine release and neural activity from ventromedial to dorsolateral striatum has been proposed to underlie reward-seeking for natural rewards and drugs of abuse (Haber et al., 2000; Haber and Knutson, 2010; Willuhn et al., 2012; Keiflin and Janak, 2015; Everitt and Robbins, 2016; Yang et al., 2018). Therefore, one might have expected an effect of DA receptor blockade in DLS on performance in our experiments. DLS dopamine, within the same anatomical location tested here, does support high levels of lever press responding for cocaine maintained by presentations of a conditioned reinforcer on a second-order schedule, an effect not observed for DMS (Vanderschuren et al., 2005; Belin and Everitt, 2008; Murray et al., 2012; Spoelder et al., 2017; Giuliano et al., 2019). This DLS dependence requires extended training, and our training was relatively brief in comparison. The nature of the instrumental sequence tasks used in the present studies may also lead to greater DMS as opposed to DLS dependence, as discussed below. Of note, Van Elzelingen and colleagues (2022) examined dopamine release using fast-scan cyclic voltammetry in striatal subregions during acquisition of a two-lever response sequence and found no evidence of ventral-to-dorsal shifts in these dopamine signals across ten weeks of training (van Elzelingen et al., 2022).

Our findings with GABAergic inactivation also suggest that behavioral control in our tasks did not extend to the DLS. Different striatal subregions are well known to mediate distinct process of instrumental behavior with the DMS mediating actions guided by representations of the reward to be earned and the DLS controlling habitual actions (Yin et al., 2004, 2005, 2006; Balleine et al., 2007; Everitt and Robbins, 2013; Corbit and Janak, 2016; Lerner, 2020), both of these tested under extinction conditions. Despite this, inhibition of neural activity in these regions has minimal or no impact when rats are allowed to complete actions that are rewarded, typically measured as responses at a single operandum (Corbit and Janak, 2007; Balleine, 2011; Vandaele et al., 2019). Here we tested the impact of a temporary lesion of these regions via pharmacological inactivation during rewarded performance of a more complicated sequence of three actions at three distinct levers and found that NAc and DMS, but not DLS, are essential for maintaining sequence performance, by controlling appropriate timing, limiting non-essential responding, and preventing serial order errors. Interestingly, inactivation of both the NAC and DMS induced errors, but only in the uncued version of the task, whereas the other performance factors were altered in both the cued and uncued task albeit with perhaps greater effects in the uncued version. These results are in contrast to findings that DLS lesion/activity suppression impairs the acquisition of a 2-response sequence across two levers, with explicit punishment for incorrect lever presses, to earn reward (Yin, 2010; Rothwell et al., 2015; Garr and Delamater, 2020), even in a case of goal-directed control (Garr and Delamater, 2020). The behavioral sequence trained here required locomotion from one to the other side of the chamber, as opposed to alternation between two adjacent levers, with the former situation perhaps more cognitively demanding, thereby relying on the DMS. It also remains possible that we failed to train rats extensively enough to result in potential recruitment of the DLS, although it is difficult to a priori determine such an appropriate amount of training (Vandaele et al., 2017), and Garr and Delamater (2020) find engagement of DLS is important for normal sequence learning and performance relatively early in training. Interestingly, explicit cuing of responses can accelerate the development of habits while other evidence suggests an increasing degree of goal-directed control when rats perform action sequences (Vandaele et al., 2017; Garr and Delamater, 2019), collectively suggesting more goal-directed control in the uncued variant of our task and potential for habit guiding cued sequence performance. Future investigations of the goal-directed vs habitual control in concert with manipulations of neural activity and dopamine release in these striatal subdivisions will be critical for identifying the psychological processes underlying action sequence performance in our procedure (Balleine, 2011; Dezfouli and Balleine, 2012; Garr, 2019; Lerner, 2020; Vandaele and Ahmed, 2021).

The ventral and dorsal striatum can be organized into distinct components of a system for executing precise actions versus the generation and memory for the steps necessary to procure reward, with the former susceptible to habitual control. Such an actor-critic dissociation may map onto both the activity of ventral and dorsal striatum and in turn the distinct dopamine projections from the ventral tegmental area and substantia nigra to these striatal subregions (Houk et al., 1995; Joel et al., 2002; Takahashi et al., 2008; Bornstein and Daw, 2011; Sutton and Barto, 2018). In this view, the accumbens core and its ventral tegmental dopaminergic input critique and refine the generation of motivated behavior, an effect that would become evident when either cues or a series of distinct movements are needed to energize and maintain reward pursuit (Bornstein and Daw, 2011; van der Meer and Redish, 2011; Hamid et al., 2016). In contrast, DLS and its dopaminergic input from the substantia nigra would selectively contribute to the precise motoric structure of action execution which would be prominent when repeated responses on a single operanda are required and stereotypy is favored (van der Meer et al., 2010; Dhawale et al., 2021). The use of a self-paced sequence of actions across three distinct operanda potentially promoted behavioral variability that prevented such nigrostriatal recruitment. These findings are also in line with our recent evidence that rats working for optogenetic activation of ventral tegmental area mesolimbic dopamine neurons, but not substantia nigra dopamine neurons, readily acquire an action sequence and exhibit flexibility in their pursuit of optogenetic stimulation in the absence of tangible reward (Fraser et al., 2022). Our results support such an actor-critic framework, with DMS having a mixed role between these two distinct subregions, but ultimately such a hypothesis requires more precise methods such as optogenetics to temporally manipulate neural activity coincident with action execution (Rothwell et al., 2015; Geddes et al., 2018; Hollon et al., 2021).

Overall we find a dissociation in the contributions striatal subregions, including their dopaminergic activity, to action sequence performance. Dopamine release in the NAc and DMS is essential for motivation to perform action sequences, with core manipulations having a greater overall impact on behavior. However, when neural activity in either the NAc or DMS is pharmacologically suppressed, other (possibly striatal) regions can apparently maintain appropriate levels of motivation. Yet, suppression of activity in the NAc and DMS does reveal that each subregion contributes to accurate sequence execution. DLS manipulations had little effect on motivation or other performance measures. These results support the notion that there exist distinct processes supported by striatal subregions and their midbrain dopamine projections that extend across Pavlovian and operant behaviors (Everitt and Robbins, 2013; Fraser and Janak, 2017; Saunders et al., 2018; Keiflin et al., 2019). Given the striatum is the primary target of these dopaminergic projections and that drugs of abuse evoke profound plasticity within striatal projection neurons (Lüscher et al., 2020; Lüscher and Janak, 2021), resolving the contribution of individual striatal subregions to precise aspects reward-seeking will be critical for developing targeted therapies for the treatment of substance abuse disorders.

## METHODS

### Subjects

Male and female Long-Evans rats (initial n=52; 24 female) were purchased from Envigo (Frederick, MD, USA) and were approximately 60 d old on arrival. Rats were single-housed in a climate-controlled facility on 12:12 light:dark cycle (lights on at 07:00). All experiments took place during the light cycle. Rats were mildly food-restricted (95% of free-feeding body weight, fed after daily behavioral sessions) starting two to three days before the beginning of behavioral experiments and maintained on food-restriction throughout all training and testing. Water was always available ad libitum. All experimental procedures were approved by the Animal Care and Use Committee of Johns Hopkins University.

### Surgery

At least one week after arrival in the animal facility, rats were anesthetized with 5% isoflurane. Once anesthesia was induced, rats were maintained at 1-2% isoflurane and placed in a stereotax for the implantation of cranial cannula. After cleaning the scalp with ethanol and Betadine surgical scrub an incision in the scalp was made to expose the skull surface. 22-gauge cannula (Plastics One) were implanted bilaterally targeting either the nucleus accumbens core (NAc; AP: +1.8 ML: ±1.5 DV: -6.0), dorsomedial striatum (DMS; AP: +1.2 ML: ±1.5 DV: -3.6), or dorsolateral striatum (DLS; AP: +1.2 ML: ±3.4 DV: -3.6). Cannula were secured to the skull with 3-4 skull screws and dental cement. Rats received cefazolin (70 mg/kg subcutaneuous) as antibiotic and carprofen (5 mg/kg subcutaneous) as painkiller immediately following surgery. At all times other than during infusions stainless steel ‘dummy’ stylets were kept in cannula that were flush with the base of the metal tubing. Rats were allowed at least one week to recover prior to the start of behavioral training.

### Behavioral Apparatus

Training and testing took place in 12 MedAssociates operant chambers housed in sound-attenuating cabinets. On one wall a recessed port equipped with a pellet dispenser containing 45-mg banana flavored grain pellets (F0059; BioServ) was flanked on either side by two retractable levers. On the opposite wall, directly across from the reward-delivery port, was a third retractable lever and a white houselight was mounted above this lever adjacent to the ceiling. When extended each lever required an approximate 10 g force to record a response. Each sound-attenuating cabinet was equipped with a fan which provided background noise. A computer running MED-PC software controlled the behavioral equipment and recorded lever presses, food cup entries, their associated timestamps, and the length of the session.

### 3-Lever Sequence Task

Rats were shaped to complete a series of three lever presses at three distinct locations to earn reward. There was one behavioral session per day. First, rats were trained to collect rewards from the port in a single session in which 100 pellets were delivered at random (10-80 s range). They were then allowed to press the lever directly to the left of the port to earn reward with each press leading to the delivery of a single pellet for 3 days. Then rats were trained to press the lever to the right of the port, the first press of which led to the insertion the lever to the left of the port, upon which the first press led to pellet delivery and retraction of the left lever. Rats were trained in this way for 3 days. Finally, rats were trained to press the lever in the rear of the chamber (the distal lever, LD), upon which the first press resulted in the insertion of the right lever (the proximal lever, LP), which the first press upon resulted in insertion of the left lever (the terminal lever, LT) which when pressed led to pellet delivery and the retraction of the left and right lever. Rats were trained in this way for 3 days prior to the first series of tests on this cued action sequence.

Following the first tests, rats were then trained for 3 days in which the first press on the rear lever resulted in the simultaneous insertion of the left and right levers. Rats then had to press the right then the left lever to earn reward coincident with the retraction of the left and right levers. For the final version, rats were allowed continuous access to all 3 levers with only the requirement to press the rear, then right, then left lever to earn reward. Training proceeded in this manner for 3 days. Rats were tested once again for the effects of dopamine antagonism and reversible inactivation on this uncued version of the task.

There were never any timeouts or punishments associated with incorrect lever presses nor did an incorrect or extra press reset the sequence. Sequences were reset only when rats entered the reward port following pellet delivery (e.g. extra presses or sequence completions following pellet delivery would not earn additional pellets until the rat entered the port). Pellet delivery was accompanied by an audible click of the dispenser and a brief 300 ms darkening of the chamber by exterminating the houselight. Sessions continued until 100 rewards were earned or 240 minutes had elapsed.

### Drug Solutions

The general dopamine receptor antagonist flupenthixol dihydrochloride (Tocris) was prepared at a concentration of 100 μM in sterile 0.9% saline and frozen in aliquots that were thawed fresh immediately prior to use. The GABAB agonist baclofen hydrochloride (Sigma) and the GABAA agonist muscimol hydrobromide (Sigma) were mixed immediately prior to use from stock concentrations to a final cocktail of 1 mM baclofen and 0.1 mM muscimol in sterile 0.9% saline.

### Intracranial Infusions

Infusions of drugs were made through 28-gauge injectors (Plastics One) connected to a 5 μL gastight Hamilton Syringe in a Harvard Instruments motorized pump via polyethylene tubing. In all cases infusers extended 1 mm beyond the base of the cannula for final infusion coordinates: NAc DV: -7.0, DMS DV: -4.6, DLS DV: -4.6. Rats were restrained by the experimenter, dummy stylets removed, injectors inserted, and 300 nL of solution was infused over the course of 1 minute with an additional minute post-infusion prior to the removal of the injector and replacement of the dummy cannula. Rats were left undisturbed following infusion completion in their home cages for 15-25 minutes prior to the start of the behavioral session. Rats were accustomed to the infusion procedure in the 3-4 days leading to the first test, and the patency of each cannula was confirmed by piercing the brain tissue with an injector the day prior to testing. Rats were tested both on the cued 3-lever task and on the uncued task variant. Each rat received 6 infusions total. The order of drug infusions (saline, flupenthixol, baclofen and muscimol) was randomized across each rat as well as between cued and uncued tests. Test sessions were reinforced as in training and terminated after 100 rewards were earned or 120 minutes elapsed.

### Histology

Brains were cut in 50 μm sections at -20 C in a cryostat and stored in 0.1 M NaPB. Sections were then mounted onto Fisher SuperFrost PLUS slides, dried, stained with cresyl violet (FD Neurotechnologies) and coverslipped with Permount (Fisher Scientific). Slides were examined under a light microscope and the location of the infuser tip was mapped onto the appropriate level of a rat brain atlas (Paxinos and Watson, 2007). Two rats were sacrificed between the cued tests and uncued tests (1 from NAc and 1 from DLS groups, respectively). Final group sizes following exclusions were: NAc (n=15; 9 male, 6 female), DMS (n=11; 5 male, 6 female), and DLS (n=12; 5 male, 7 female).

### Statistics and Data Analysis

Data were processed in Excel (Microsoft) and Neuroexplorer (Nex Technologies), analyzed with SPSS (IBM), and visualized Prism 9 (GraphPad). We calculated sequence completion length as the time from the first press on LD to the first press on LT which resulted in reward delivery. We also separately analyzed the time between the first LD and LP press and the first LP and LT press for each sequence. Sequence times and interpress times were log-transformed to adjust for positive skew. Task efficiency was defined as the number of correct lever presses relative to the total number of lever presses. We defined perseverative errors on each lever as extra presses that occurred within that segment of sequence completion (e.g. perseverative LD presses were those that occurred between the first LD press and the subsequent LP press) and sequence errors reflected responses on a lever that would represent either a jump forward (e.g. LT responses in the segment between the first LD and subsequent LP press) or a regression in the sequence. Training data were analyzed with generalized linear models with a first-order antedependence covariance structure and any relevant posthoc comparisons were made with Bonferroni’s method. Testing data were analyzed with repeated measures ANOVA with post-hoc comparisons made with Dunnett’s test to compare each drug condition relative to saline control. For all analyses α=0.05.

## Notes

### Competing Interest Statement

The authors have declared no competing interest.

